# Verification and reproducible curation of the BioModels repository

**DOI:** 10.1101/2025.01.16.633337

**Authors:** Lucian Smith, Rahuman S. Malik-Sheriff, Tung V. N. Nguyen, Henning Hermjakob, Jonathan Karr, Bilal Shaikh, Logan Drescher, Ion I. Moraru, James C. Schaff, Eran Agmon, Alexander A. Patrie, Michael L. Blinov, Joseph L. Hellerstein, Elebeoba E. May, David P. Nickerson, John H. Gennari, Herbert M. Sauro

**Author notes:** **Author contributions** - LS: Writing, Conceptualization, Data Curation, Investigation, Methodology, Project Administration, Software, Validation. - RMS: Reading, Writing, Data Curation, Methodology - TN: Reading, Data Curation, Methodology - HH: Reading - JK: Conceptualization, Data Curation, Investigation, Methodology, Software. - BS: Software - LD: Software - IIM: Reading, Conceptualization, Funding - JCS: Software, Methodology - EA: Reading, Writing - AAP: Software - MLB: Reading, Writing - JH: Writing, Methodology - EM: Reading, Writing - DPN: Reading, Writing, Methodology - JG: Reading, Writing, Methodology - HMS: Reading, Writing, Funding.

## Abstract

The BioModels Repository contains over 1000 manually curated mechanistic models from published literature, most often encoded in the Systems Biology Markup Language (SBML). This community-based standard formally specifies each model, but does not describe the computational experimental conditions to run a simulation. Therefore, it can be challenging to reproduce any figure or result from a publication with an SBML model alone. The Simulation Experiment Description Markup Language (SED-ML) provides a solution: a standard way to specify exactly how to run an experiment corresponding to a specific figure or result. BioModels was established years before SED-ML, and both systems evolved over time, both in content and acceptance. Hence, only about half of the entries in BioModels contained SED-ML files, and these files reflected the version of SED-ML that was available at the time. Additionally, almost all of these SED-ML files had at least one minor mistake that made them invalid. To make these models and their results more reproducible, we report here on our work updating, correcting and generating new SED-ML files for 1055 curated mechanistic models in BioModels. In addition, because SED-ML is implementation-independent, it can be used for *verification*, demonstrating that results hold across multiple simulation engines. We tested, corrected, and improved over 450 existing SED-ML files in the BioModels database, and created basic files for the rest of the entries. Then, we used a wrapper architecture for interpreting SED-ML, and report verification results across five different ODE-based biosimulation engines, after further improving the models, the wrappers, and the engines themselves. Our work with SED-ML and the BioModels collection aims to improve the utility of these models by making them more reproducible and credible. Improved reproducibility means these models are now even more fit for re-use, such as in new investigations and as components of multiscale models.

**Author summary:** Reproducing computationally-derived scientific results seems like it should be straightforward, but is often elusive. Code is lost, file formats change, and knowledge of what was done is only partially recorded and/or forgotten. Model repositories such as BioModels address this failing in the Systems Biology domain by encoding models in a standard format that can reproduce a figure from the paper from which it was drawn. Here, we delved into the BioModels repository to create and correct the instructions on what to do with every curated model, and then tested those instructions on a variety of simulation platforms, allowing us to find and correct issues with the platforms and the simulators themselves. Not only did this improve the BioModels repository, but also improved the infrastructure necessary to run these verification comparisons in the future, and improved the fitness of the models for re-use by other researchers.

## Introduction

The reproducibility of scientific research is one of the cornerstones of the scientific method. It may therefore be surprising to learn that the reproducibility of scientific work has been of some recent concern [1]. Numerous articles [2] and reports [3] have been published that discuss reproducibility challenges and possible remedies. These challenges can apply to both wet lab experimentation as well as computational studies. As an example, a recent survey by the BioModels team [4] confirmed previous anecdotal evidence that a large proportion (over 50%) of published computational models of physiological processes were essentially irreproducible based on the information provided in the manuscript.

Here, we focus on the reproducibility and verification of computational models published by the systems biology and physiology communities. These are simulation studies based on a proposed mechanistic model of some biological process. For the purpose of this paper, we consider reproducibility to mean that the scientific claims of the paper can be verified by reproducing the execution of the computational model, and verification to mean that separate simulation engines produce the same results when running the same computational experiment. Both are key to model fitness for re-use and expansion in new contexts, as all science strives to be.

In theory, the results from computational models should be easily reproducible, as the mathematics involved is well understood, and the calculations should be the same regardless of the particular software or operating system used. However, modeling has been no more immune to reproducibility failures than any other branch of science [1], and attempts to reproduce even well-known models in the field have encountered a wide variety of problems [5].

In this paper, we will examine the reproducibility and verification (defined below) of the large corpus of computational models held at the BioModels repository [6]. Before presenting details of our methods and results, we provide context and background about standards development in the modeling community and the establishment of curated collections of models, such as BioModels.

### Modeling Standards

#### SBML

In the last 15 years, the reuse and long-term storage of computational models generated by the biomedical research community has been greatly improved by the use of community modeling standards [4, 7]. One of the more important such standards is the Systems Biology Markup Language (SBML) [8]. SBML is a standardized, machine-readable format for representing computational models of biological processes. The emergence of SBML stimulated the development of repositories for storing models. One of the most well-known of these is BioModels [6] but others exist, such as BiGG [9], KBase [10], and Model SEED [11]. The models stored at BioModels are mostly kinetic models that have been obtained from the literature, and the collection includes metabolic, protein signaling, and gene regulatory models. BioModels has grown to include almost 1100 curated models (as of May 2025) collected from peer reviewed articles. These models have been *manually* curated by staff at the European Bioinformatics Institute (EMBL-EBI) to ensure that they reproduce published results. Although EMBL-EBI established a consistent protocol for their curation, the process was manual, and no computational method or log was kept about how curators executed the models or matched published results with reproduced ones.

The BioModels repository serves as a critically important resource for the systems biology community. Using one of the many simulation tools [12] that support SBML, models downloaded from BioModels can be simulated with the expectation that results will match what was reported in the original publication. In this work, this expectation was exhaustively tested, allowing us to illuminate and correct several gaps between the expectation and reality.

#### SED-ML

SBML only describes the model; it provides no information on how the results were generated in the associated published article. The community recognized this as a significant failing because it meant that other researchers would not be able to easily reproduce the results of a published article. As a result, in addition to SBML, a second community standard, the Simulation Experiment Description Markup Language (SED-ML) was developed [13, 14]. The purpose of SED-ML is to describe how the simulation results in a given research article are generated.

SED-ML aims to capture the process of loading a model, potentially changing model values, running a simulation experiment such as a time-course simulation or parameter scan, and collecting the results as tables of data (reports) or plots. Importantly, none of these steps is connected to any particular simulation application. Thus, SED-ML provides a declarative, software-independent language for describing a simulation experiment.

BioModels did include SED-ML files for about half of the curated models; however, *caveat emptor* : they existed and were potentially useful, but had not been checked. In this work, each file’s usefulness has now been improved and clearly defined. Figure 1 shows a schematic of how reproducibility is supported by these two standards.

**Fig 1.**
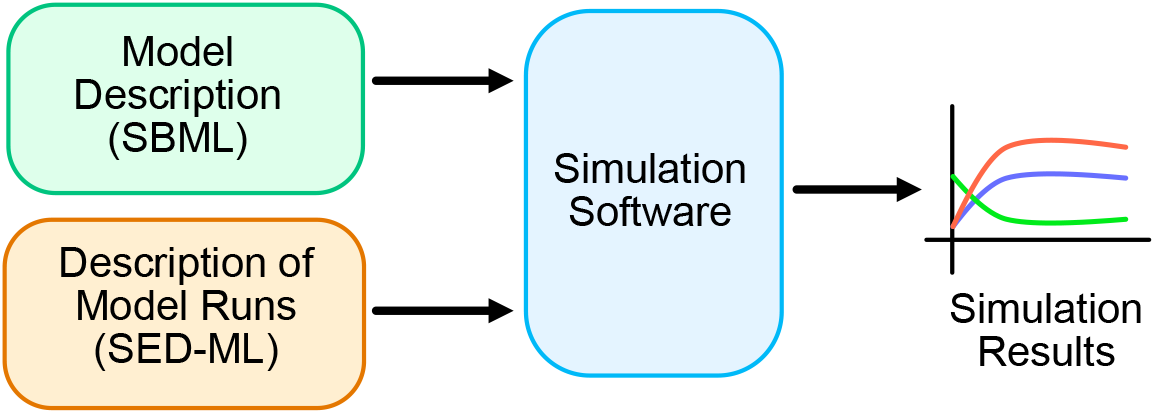
Combining SBML and SED-ML to generate simulation results.

#### OMEX

To keep the SBML, SED-ML, and other files together, they can be stored in Open Model EXchange (OMEX) files, a format standardized by the biomodeling community as a standard way of collecting model experiment files together [15]. This format is essentially a ZIP file of the collected files, plus a manifest file listing and describing each file. In 2017, the BioModels database was updated to allow each entry’s collection of files to be downloaded as an OMEX file, with the manifest file noting which file was the canonical model file for that entry.

### The BioModels Protocol for Curating Models

The EBI-EMBL protocol for curating a model in the BioModels collection begins with encoding the model into a standard format, typically the Systems Biology Markup Language (SBML). The curator then selects a figure from the publication and attempts to reproduce this figure by creating a simulation experiment using simulation software. Since BioModels is primarily an SBML repository, the simulator must be able to interpret the SBML. However, many such simulators exist, such as COPASI [16], Tellurium [17] or VCell [18]. The curator will then provide the output figure from the software engine, and a brief description of the curation work (e.g., (from BioModel 836), *“Two figures were drawn from the model based on the data provided in the paper. The values of panic intensity and protection were changed to 0*.*1 and 0*.*1 by trial and error*.*”*). Finally, model elements are annotated, following an established procedure for annotation based on MIRIAM guidelines [19] (the Minimal Information Requested In the Annotation of biochemical Models). Once the curation is complete, all other required and auxiliary files are placed into an archive for that entry.

Although this process establishes that a set of results can be reproduced, there are some shortcomings. Curation often involves some guessing on the part of the curator, which can make the process non-repeatable if those choices are not recorded. The curation results may depend on which figure is being reproduced, the simulator settings, the time steps, initial conditions, and other considerations that must be inferred from the published article. This kind of information can be stored in SED-ML [13] files alongside the SBML model.

### Verification of Models

A number of organizations have discussed the need for improving the credibility of computational models [20–22]. The credibility of a model is strengthened if it can be verified, validated, and its uncertainty measured or quantified. These three elements are often abbreviated VVUQ, for verification, validation, and uncertainty quantification.

In this study, we focus on the verification of models described by SBML and SED-ML in the BioModels collection. We use the definition of verification from the Los Alamos National Laboratory white paper [23], which defines verification as ‘… the process of determining that a model implementation accurately represents the developer’s conceptual description of the model and the solution to the model.’ It is important to note that verification does not consider whether the model is a faithful representation of the biological system under study; that would be the concern of validation. Verification is used to determine whether the numerical algorithms and model description have been implemented correctly in software. Until the development of standards such as SBML and SED-ML, automatic verification was not possible for biosimulation models.

### BioSimulators

The Biosimulators site is a repository of simulation applications that support modeling standards (biosimulators.org). Each of these simulation applications is packaged as an interchangeable “biosimulator” for running simulation experiments encoded in SED-ML. These biosimulators collectively span a broad range of simulation capabilities with enough overlap to demonstrate model reproducibility and to cross-validate predictions across multiple simulators. Each biosimulator simulation tool computes SED-ML simulation experiments for a subset of modeling formats (SBML, NeuroML, CellML), frameworks (logical, kinetic) and simulation algorithms (FBA, SSA) and are available at biosimulators.org/simulators.

The biosimulations.org website allows users to run simulation experiments specified in SED-ML, and view results and plots, and hosts a searchable collection of published simulation experiment projects, many of which were created from BioModels.net database models. APIs are provided for validating the correctness of SED-ML, models and metadata documents (combine.api.biosimulations.org) and for running uploaded simulation experiments and downloading the simulation results (api.biosimulations.org). A dedicated API service compares the results of multiple versioned biosimulators when exercised with the same simulation experiment https://biosim.biosimulations.org/docs

### Summary of Results

In this paper, we have two main results to report. First, existing SED-ML files from curated BioModels were tested for the first time, and numerous errors were corrected. If a model had no SED-ML, we created a ‘template’ SED-ML file with a generic simulation experiment so that the model’s behavior could be verified across different simulators. Next, with SED-ML in hand, we report on preliminary results from a verification study of all curated models across five different biosimulation platforms. This verification study allowed us to further improve the SED-ML and SBML files, as well as the simulators themselves.

To carry out this work, we took a snapshot in time from June of 2024 of the curated branch of BioModels, working with a copy of the entire curated branch of 1073 models, 1055 of which were ODE models. Of these 1055 models, we were able to create SED-ML files that produced the same results for at least two biosimulation engines in 932 cases (88%). To our knowledge, this work is the first systematic exploration of different simulators’ interpretations of SED-ML constructs. Although these early results are not perfect, the use of SED-ML provides a way forward. These formal specifications can now be used to investigate the differences discovered across biosimulation engines, verifying not only the models themselves but the simulation engines as well.

### Methodology

Although there are many ways to improve the reproducibility of systems biology modeling, here we focus on the ability of researchers to replicate published simulation results and then to verify them using different biosimulation engines. As Figure 2 shows, as a first step, model repositories such as BioModels should include SED-ML to describe the steps needed to replicate the results. We have created a wrapper architecture at BioSimulators.org [24] where this information can then be interpreted and sent to different biosimulation engines. The majority of models in the curated BioModels collection are ODE models (1055 of 1073); therefore, we have selected five well-tested biosimulation engines, and built and tested wrappers for these: COPASI [16], Tellurium [17], VCell [25], PySCeS [26] and Amici [27]. As we describe, after many fixes and updates, many curated models (but not all) could indeed be verified across more than one of these biosimulation engines.

**Fig 2.**
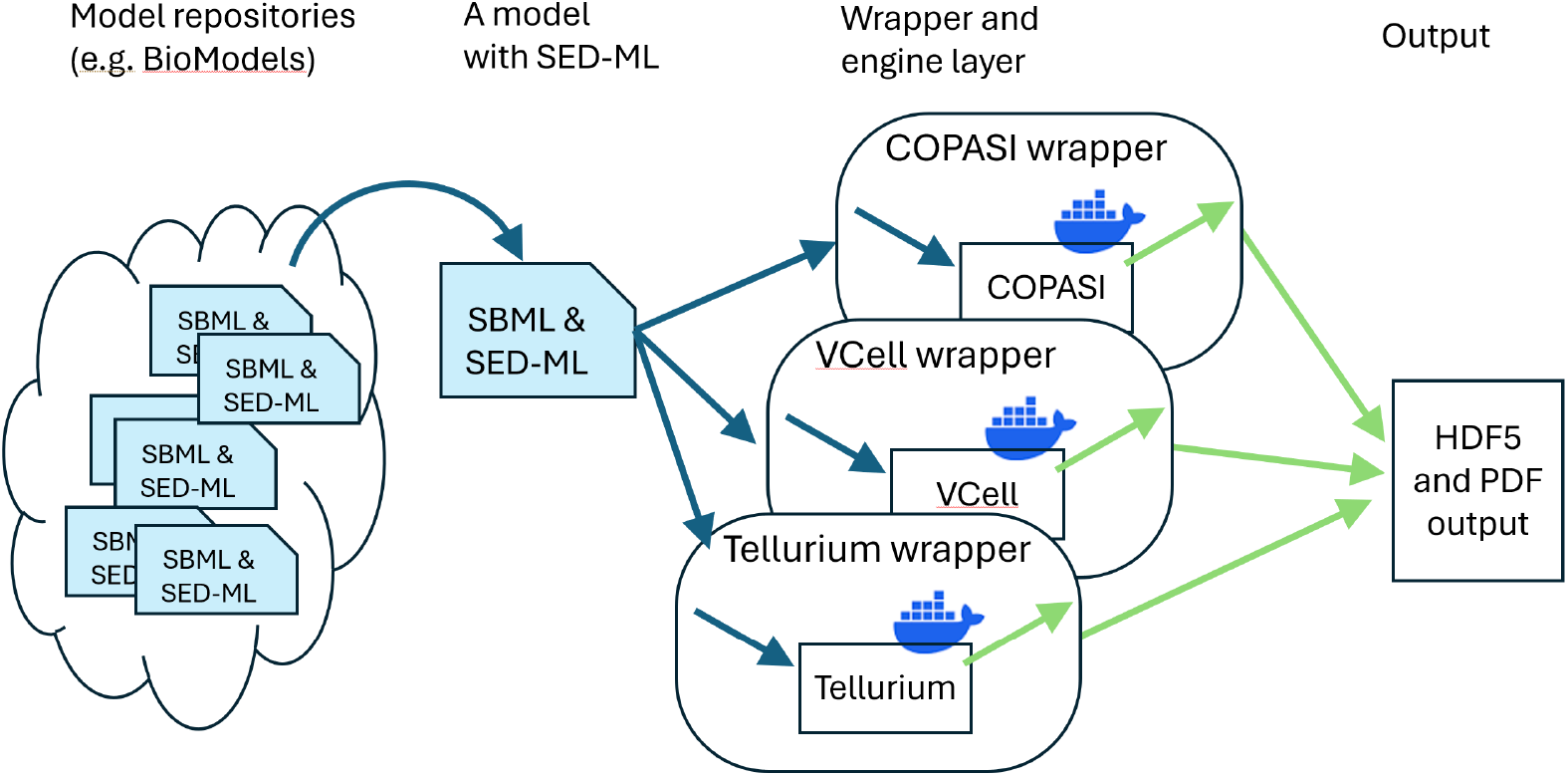
SED-ML and wrappers that allow researchers to replicate results across multiple simulators. See Shaikh et al. [24] for more details about the biosimulation engine wrapper architecture.

### SED-ML Wrapper Development

Most biosimulation software engines are developed independently, each with its own unique interfaces. While many offer APIs (Application Programming Interfaces) for accessing their capabilities, these APIs differ from one engine to another, requiring customized handling of input and output data to match each engine’s specific API. Our solution, described previously [24] and illustrated in Figure 2, is a “wrapper” architecture: each wrapper translates SED-ML instructions to API calls for the respective biosimulation engine, all within Docker containers to ensure platform independence and stability over time. By using this approach, SED-ML serves as a *lingua franca*, enabling the same procedure to be executed across different wrapped biosimulation engines. This allows researchers to precisely verify results using standardized inputs and outputs across multiple engines.

Each wrapper takes an OMEX file containing at least one SED-ML and one SBML file as input, translates the SED-ML instructions into simulator-specific commands, collects the output of any simulations from those simulators, and exports all outputs. Tabular data is exported as HDF5 files [28] (a format similar to CSV, but allowing multi-dimensional data and annotations), and figures are exported as PDF files. While non-semantic differences in figure output prevent the automatic comparison of figure data, the numbers inside the HDF5 files can be compared directly. In this way, we can verify whether the SED-ML files will produce the same output on multiple biosimulation engines.

In the process of developing the wrappers, we discovered a number of SED-ML constructs that were ill-defined in the specification (which at that time was Level 1 Version 3, released October 2017 [29]). We coordinated with the SED-ML community and the SED-ML Editors in particular to clarify and update the specification. As a result, the SED-ML Level 1 Version 4 specification was released in September 2021 [30] (with some members of our team joining as SED-ML editors), which included over 100 changes due to issues discovered during this process.

### BioModels secondary file validation

At the core of our process was a collection of Python programs, available at https://github.com/sys-bio/temp-biomodels, in concert with GitHub actions that would run the program over all 1073 BioModel curated entries. This project performed many general curation and validation steps, which will be directly applicable to future validation and curation efforts, as well as a large number of subroutines to fix problems with particular files.

The first step of the program is to simply validate all the files. The BioModels entries contained a total of 11,890 files of 24 different types, from SED-ML files to original COPASI files to images and PDF documents. At EMBL-EBI, the BioModels curation process (described above) has generally been focused on the SBML model file, with all ancillary files treated as secondary to the SBML file. These secondary files had never before been systematically checked for accuracy, and our first goal for this project was to ensure that the files were at least valid. The program performed some basic maintenance: 40 files of size zero were discovered and removed; filenames with illegal characters were renamed; filenames of one type were renamed to a consistent scheme (‘.sedml’; ‘.ode’); a few filename typos were fixed. We found validators for all 24 file types, and applied them to every file in the archives, finding several that were invalid. We then examined the invalid files to see if the problems could be fixed. If so, additional steps were added to correct those individual files, and if not, the files were deleted. Other problems were also discovered and corrected, such as auto-generated PDF files containing only an error message.

Several files had been auto-generated by a variety of translation programs that themselves came and went over the 20 years that BioModels has been active. We were able to track down the latest versions of some of those translators, and regenerated those files. This ensured that updates to the translation algorithms developed over the last decade plus would be incorporated into the latest versions of the files. These translators were additionally used to create new files for entries that previously had not contained them.

Many SBML files were present in two forms: one with annotations in URN form, and another with annotations in URL form. This reflected the diversity of opinion in the SBML community when BioModels was first created as to which format was more useful. In the intervening years, that debate has been concluded, with the URL form gaining universal acceptance. Thus, the legacy URN-formatted SBML files were removed as well.

We also validated the final annotation step of curation by ensuring that every annotation URL in the SBML file pointed to an actual entry in an existing ontology. A variety of errors were caught by this check, including simple typos of the name of the ontology, or leaving out parts of the required format for the reference. Of the over 208,000 annotation references, approximately 70 errors were discovered and manually corrected, consulting the original paper for references when necessary.

### BioModels SED-ML updates

Our effort to generally clean up the entries paled in comparison to our effort to update and test the SED-ML files. In a perfectly-curated entry, there would be a SED-ML file present, with instructions for how to reproduce a figure from the curated paper, encoding in a reproducible way the process the original curator of the model undertook when creating and testing the model. Barring that, there should be a SED-ML present that describes some sort of simulation of the model that could be replicated across multiple simulators.

We started with the two types of files that were present: SED-ML files and COPASI files. Helpfully, the current version of COPASI (4.43) is backwards compatible with older COPASI files, meaning that we could create SED-ML for many entries that had been created before SED-ML even existed. When both a COPASI model file and a previously-exported SED-ML file were present for the same model, we hand-compared the existing file to one recreated from the COPASI file. In most cases, the files were nearly identical or the newly recreated files were more precise and detailed. In three cases, the existing SED-ML file had more details than the re-generated version. We hypothesized this was due to the SED-ML having been exported from a working COPASI curation session which was not subsequently stored as a COPASI file. The results of these hand comparisons were encoded into our Python curation program.

For the 579 entries with no SED-ML nor a COPASI file that could be used to generate SED-ML, we created a ‘template’ SED-ML file, encoding a simple time course experiment into it. The model is loaded, a timecourse simulation is performed for ten time units, and finally, the levels of all variable species are exported, both as a table of values per time point, and as a plot. Of course, these files do not match any figures from the original publication, but can still be used for verification purposes: if two simulators produce the same output from this template SED-ML, it shows that the model itself is robust to interpretation. It is also much easier to edit an existing SED-ML file than to create one from scratch, making future efforts to reproduce published figures simpler [31].

### BioModels simulation testing, corrections, and upgrades

A separate GitHub repository (https://github.com/biosimulations/biosimulations-runutils) was developed to run OMEX files on our wrapped simulators using the https://biosimulations.org/ API, and collect and compare the results. Unlike the temp-biomodels repository, this project was created to be generally usable: anyone can submit an OMEX file as input and compare the resulting output across simulators. We used it as part of our workflow to simulate all 1055 BioModels ODE OMEX files on five wrapped simulators, compare results, make improvements, and repeat.

When we first attempted to verify results across multiple simulators with COPASI-generated SED-ML files, almost every single run failed due to a limitation of COPASI: every SED-ML file pointed to a ‘model.xml’ SBML file that did not exist, since COPASI only knows about its internal model, and has no way to know what filename it might have been exported as. This is true both for the SED-ML files we generated from the original COPASI files as well as the existing legacy SED-ML files found in existing BioModels entries, making all of them invalid apart from the five that had been created by hand. This reinforced the notion that SED-ML was being produced by curators as a perceived convenience to users but never checked, or this obvious deficiency would have been revealed.

So the next goal of our Python curation process was to add a fix to replace the fictitious ‘model.xml’ references with actual SBML files. When only one SBML file was present, this was trivial; when multiple SBML files were present, heuristics were developed based on file names and content similarity.

This gave us a complete, testable pipeline: SBML and SED-ML files were present that could be sent to our five wrapped simulators, and results could be compared. We could cross-verify using the native SED-ML support in Tellurium [17], separate from its wrapped version. Our process was to run a given BioModels entry across several wrapped simulators, test it in Tellurium directly, and check for error messages or empty or mismatched output. Errors would be discovered and fixed, and then we would run the entire pipeline again. This was clearly the first time anything like this had been attempted, as hundreds and hundreds of problems were discovered.

Many problems were present in the SED-ML files themselves, beyond the ‘model.xml’ issue: references were found to nonexistent model elements; simulations were defined and never used; duplicate elements were present; pointers to model elements were incorrectly formatted; simulation parameters were incorrectly applied; and many other issues were present. Other more prosaic issues were also discovered, such as requests for millions of output data points, which could bog down simulators for hours. As each issue caused a simulator run to fail, these issues were discovered, and fixes were added to the Python curation pipeline so fixed files could be further tested.

Occasionally, issues were found in the original SBML models. The most common issue we found were otherwise unused parameters which were mistakenly initialized to infinity or NaN. Models were also found with invalid SBML constructs in packages such as Layout or Render (used to store visualization information in the models). Fixes for these problems were also added to the Python curation pipeline.

In some cases, we could address issues in the simulators by adding workarounds in the wrappers. For example, Tellurium had no way to adjust SBML ‘local parameters’ on the fly, so adjustments were made to the wrapper to translate local parameters to global parameters, where they could then be adjusted. Similarly, PySCeS does not support SBML ‘initial assignments’, so we added instructions to the wrapper to translate initial assignment formulas to numerical values before asking PySCeS to simulate the model. A few of our wrapped simulators will not export constant parameters as output, so routines were added to the wrappers to add them when requested by the SED-ML.

The simulators themselves also were fixed and upgraded. The simulators worked on by our own development teams (Tellurium and VCell) were straightforward to add fixes and enhancements to, but this work also resulted in upgrades and enhancements to COPASI and PySCeS as well, as runs were found that failed on those simulators, bug reports were filed, and the development teams of those simulators produced new releases.

Other issues required more systemic solutions. As one dramatic example shows, Figure 3A and B show the original simulation results from COPASI and Tellurium (respectively) for Biomodel 1, a basic model with a template SED-ML file. The results for Tellurium are clearly incorrect, with several concentrations going negative. After experimentation and discussion with the COPASI developers, we discovered that while both COPASI and Tellurium have a setting called ‘absolute tolerance’ with a default value of 1*e*^*−*12^, COPASI then creates an absolute tolerance vector, using that value, but scaling it by the initial values of every variable in the simulation before sending that to the solver. When Tellurium was updated to allow the same technique with its own solver, it was able to replicate the COPASI results (Figure 3C). This information was then added to the SED-ML by creating a new ‘absolute tolerance adjustment factor’ setting for the simulation algorithm in the KiSAO ontology [32], which SED-ML uses to fine-tune simulation runs. The wrappers were all updated to understand and properly apply that new term, and since these results were more robust, Tellurium itself was modified to use an absolute tolerance adjustment factor by default, instead of using its old behavior.

**Fig 3.**
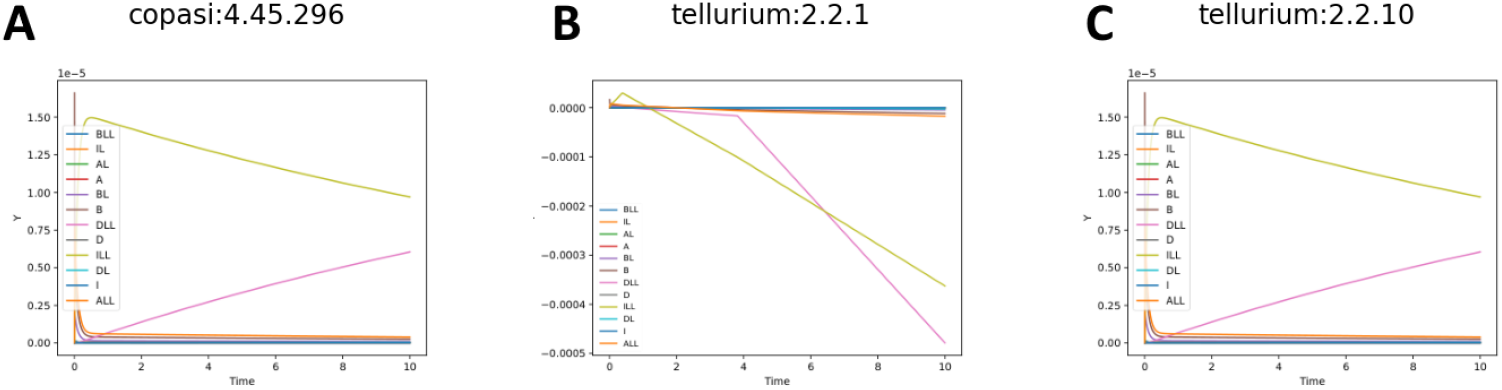
BioModels 001 simulated with Copasi (A), an older version of Tellurium (B), and the latest version of Tellurium (C).

Even this single fix has therefore improved the user experience on several fronts: model creators can encode SED-ML files more robustly with greater certainty that their results can be replicated; users of BioModel 1 (and several other models with the same phenotype) now have more varied options of simulators on which to run it; and users of Tellurium now have a more robust simulation engine for their models.

These new files were then pushed to the official BioModels repository at http://biomodels.net, where they all show up as the latest versions of the curated models. This means that executable SED-ML is now available for all of these models, each noting whether it is a ‘template’ file or one drawn from the curation process.

Moving forward, the scripts developed can be used by EMBL-EBI curators to perform more robust curation: not only can the files they collect be validated, but they can now additionally store validated SED-ML files and test them across multiple biosimulation engines on biosimulations.org.

## Results and Discussion

All 1073 curated entries retrieved from the BioModels database have been updated, with every file now validated or replaced. All 1055 ODE entries now have a new valid SED-ML file that can be used to simulate the model in some capacity: there are 579 “template” SED-ML files, and 476 SED-ML files that encode a simulation created as part of the original curation process (likely reproducing a figure from the publication).

As shown in Table 1, the great majority of these files run with the Tellurium and Copasi wrappers, and hundreds of models run successfully using our other three SBML wrappers. Tellurium has the most, simply because we focused our efforts on the wrapper for this platform since we have the most knowledge and control over this biosimulation engine. Work is ongoing to update the other wrappers to match. Figure 4 shows the distribution of successful runs and replications for both template and “full” SED-ML; i.e., SED-ML code for the 476 models from the curation process.

**Table 1.** Successful runs for each simulator of the 1055 currently curated ODE-based BioModels.

| Simulator | Successful runs |
| --- | --- |
| Tellurium wrapper | 1036 |
| Copasi (basico) wrapper | 1032 |
| VCell wrapper | 893 |
| Amici wrapper | 758 |
| PySCeS wrapper | 711 |
| At least two of the above | 1031 |

**Fig 4.**
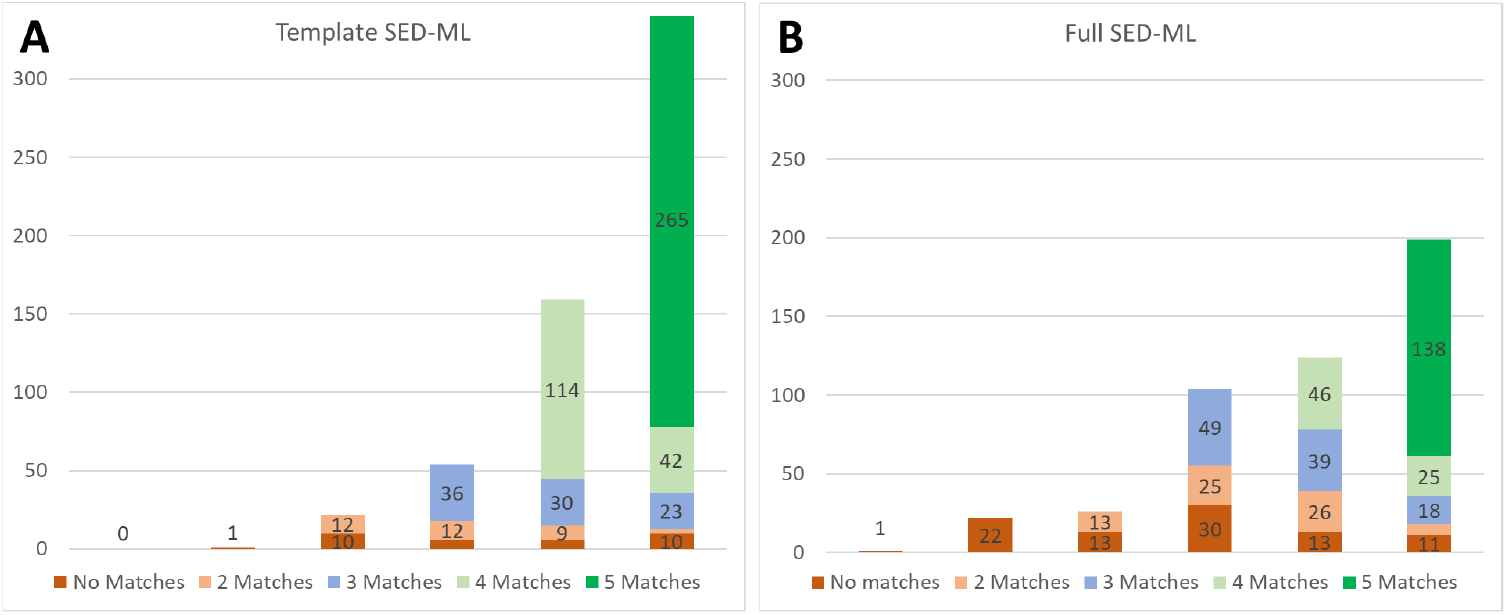
Template (A) and Full SED-ML (B) replication across five wrapped simulators.

For both parts of Figure 4, the height of the full bar indicates the number of models that successfully ran on zero (left) through five (right) of our wrapped simulators. Both the leftmost ‘zero’ and ‘one’ columns could only have no matches, as there were not two successful simulations to compare against each other. For the most part, the ‘full’ SED-ML entries did not run on as many simulators as the ‘template’ SED-ML entries, as the latter use a wider variety of SED-ML features, not all of which are supported by all the wrappers or simulators.

For all models with successful runs on at least two wrapped simulators, we defined a successful replication as having results that matched to a relative tolerance of 0.0001 and absolute tolerance scaled by the range of output values for that variable. As can be seen, only a single ‘template’ SED-ML file only successfully ran on a single simulator, and of the ‘full’ SED-ML file entries, one did not run on any of the five simulator wrappers, and 22 only ran on one. These entries were unable to be verified because two sets of results did not exist to be compared. Of the remaining entries with the relatively straightforward template SED-ML, replication was possible in 578 cases, and successfully verified in 546/578 cases (94%). For the potentially more complicated entries with full SED-ML, 453 cases could be replicated, with verification shown for 386/453 cases (85%).

Initially, our results for replicability across engines were much weaker. An earlier draft of this manuscript showed much lower success rates for simulation engines other than Tellurium. However, once these early results were circulated among the community, a broad and significant effort was invested by many institutions and research groups. Bugs were fixed and improvements were made to the SED-ML files themselves (such as fixing references or ranges), the biosimulation wrappers (such as fixing SED-ML interpretation, and improving edge case handling), the infrastructure that runs the wrappers (such as improving latency and robustness), and the simulation engines themselves (such as improving SBML interpretation and handling of simulation edge cases). Thus, an important contribution of our work has been to stimulate this type of community-based work to improve the consistency and accessibility across a range of simulation engines. This work is not finished, and is expected to continue into the future, spurring further improvements.

All of these improvements are freely available: the SED-ML files are now available at part of the BioModels entries at http://biomodels.net, the wrappers are available on github at https://github.com/orgs/biosimulators/repositories, the Docker images are available at https://biosimulators.org/simulators, the wrappers can be run online at https://biosimulations.org/runs/new, or by using our Python package https://github.com/biosimulations/biosimulations-runutils, results from verified BioModels entries (and others) can be viewed and downloaded from https://biosimulations.org/projects, and the improved simulators can all be accessed from their authors’ web sites.

Furthermore, additional improvements in consistency will now be easier to carry out. Simulators that fail to produce results from particular BioModels entries now have a test harness where improvements to the simulator or simulator wrapper can be checked as they are expanded to handle a wider variety of inputs. Cases where a simulator’s results for a given BioModel entry did not match replicated results from two or more other simulators can now be pulled out and examined: at least 30 such models exist for every single simulator in our study.

Our work also allows future BioModels curators to avoid these problems from the beginning. Using the scripts we created for this project, curators will be able to generate and test SED-ML instructions more easily, and verify results across simulators.

Recall that this study stops short of ensuring that the simulator runs actually reproduce a figure from the original publication; for many models, all we have is brief “curator comments”, so this process cannot be easily automated. We expect that most of the 476 entries (see Figure 4b) that contained SED-ML or COPASI files will indeed match a figure from the paper, as those files were originally created during curation. However, as the inclusion of a COPASI file was not a formal part of the original curation process, this is not guaranteed, so every model will need to be checked by hand. The entries with template SED-ML will not match published results, and the SED-ML would need to be modified to match the original computational experiment.

It is now also increasingly possible to usefully critique the models themselves: how reproducible are they? How sensitive are the published results to the parameter values? Can they be extended? With a system in place that makes re-simulation and verification straightforward, these questions become much easier to answer.

### Conclusions and future directions

This project demonstrated that even manually curated models can be challenging to reproduce without appropriate specifications for how to carry out the experiment. We demonstrate the utility of the SED-ML language for capturing these specifications by comprehensively creating or improving valid SED-ML files for 1055 ODE-based models in the BioModels database, all but one of which could be successfully run in at least one of our five wrapped simulation engines. For 88% of the models (928/1055), we were able to show that two separate simulators produced the same results within reasonable tolerances. Work still needs to be done to fully replicate the original curators’ work in reproducing the published results, as there is no formal record of how these models were originally curated. We have begun some of this work, and early results are encouraging [31].

Moving forward, this project demonstrates the need for improved SED-ML generation tools. Model-building environments should be able to generate SED-ML for any given experiment or execution of a model. The COPASI tool comes close to this, but it only creates SED-ML for a single model/experiment/plot combination, which isn’t sufficient for experiments involving multiple models, nor plots involving multiple experiments. In addition, it cannot export model changes (i.e. ‘in this figure, the value of n was 2, and the value of S1 was 2.42’). As an alternative approach, the Tellurium Python environment includes the phraSED-ML library, which makes it easy to create and execute a SED-ML file, but ‘creating a SED-ML file’ has to be a goal of the user, who otherwise will just use Python directly to carry out their experiments.

In general, auto-generating SED-ML will be challenging. In a GUI environment like COPASI, model changes are executed and not stored, so an experimental protocol that includes ‘change this value’ will not encode that change into the SED-ML, but will instead export a different initial model with those new values. In a scripting environment like Tellurium, common scripting behaviors like loops will commonly be encoded into the scripting environment itself, making it difficult to then export to SED-ML. Any tool that produces SED-ML but requires the user to encode these procedures separately from the user’s usual workflow must therefore offer significant benefits in exchange.

Finally, an important contribution of our work is to demonstrate the value of the wrapper architecture for biosimulation engines, and how that can support model verification. As we have shown, although many models do produce the same results across multiple engines, some models do not execute on some engines, and some models produce different results across engines. All of our scripts and wrappers are available via GitHub (https://github.com/biosimulators/), and we hope that model builders will be interested in using these to test their experiments across multiple biosimulators. In the future, we plan to build a REST API for verifying any model coded in SBML (and/or OMEX archives with SBML) across multiple biosimulation engines.

Although the BioModels database has carried out ground-breaking work developing a curated model repository, until now, the database has not included enough information to fully replicate published results. Here, we have made significant inroads towards solving this problem by providing valid SED-ML files for all models, as well as SED-ML files involved in curation for almost half of the collection. Moving forward, additional work on improved wrappers is needed, and model development environments should be designed to facilitate the auto-generation of SED-ML. Without strong tool support and systematic checking of results, it is too easy for errors or omissions to accumulate, which ultimately makes published results non-reproducible. To ensure reproducible results, we strongly encourage authors to submit their models, accompanied by correct SED-ML files, to BioModels or comparable resources as part of the publication process. With the use of SED-ML, we have demonstrated the ability to *verify* published results across multiple biosimulation engines. This increase in reproducibility has in turn increased the value of the repository to modelers going forward. New models will be directly comparable to existing ones using the same protocols, and the existing models can be expanded and used in new contexts with greater assurance of their utility.

## Acknowledgments

This work would not have been possible without the years of work from over 200 BioModels curators and modelers who submitted their work to the BioModels repository.

The Center for Reproducible Biomedical Modeling is funded by the NIH (grant ID P41EB023912). IIM acknowledges additional funding from NIH grant R24GM137787. RMS, TN and HH acknowledge EMBL Core Funding. This work has also received funding from the Innovative Medicines Initiative 2 Joint Undertaking under grant agreement number 116030 (TransQST). This Joint Undertaking receives support from the European Union’s Horizon 2020 Research and Innovation Programme and European Federation of Pharmaceutical Industries and Associations (EFPIA).

## Notes

### Competing Interest Statement

The authors have declared no competing interest.

### Summary of Updates

The new version of the paper has been greatly expanded, from nine and a half pages to thirteen. Every section has received major updates, these include: * The Introduction section now better sets up the state of the art in the field, the motivation, and how our work contributes to and extends that. * The Methods section was expanded to include many more details, including a new figure illustrating a typical issue that was addressed. * The Results and Discussion section was expanded to emphasize how specifically this work contributes to the field.

